# *Marchantia* liverworts as a *proxy* to plants basal microbiomes

**DOI:** 10.1101/103861

**Authors:** Luis D. Alcaraz, Mariana Peimbert, Ana E. Dorantes-Acosta, John L. Bowman, Mario A. Arteaga-Vázquez

## Abstract

Microbiomes influence plant development, establishment, nutrient acquisition, pathogen defense, and the myriad of roles that ultimately impacts plant health. Plants microbiome are shaped through interactions between the microbes (ranging from cooperative functions to chemical warfare) and a selection process entailed by the host plants that distinguishes between pathogens, commensals, symbionts and transient bacteria. The soil is a primary source for microbes colonizing plants, along with other environmental sources including rain and interactions with other organisms. In this work, we explore the microbiomes through massive sequencing of the 16S rRNA gene in the eldest terrestrial plants: *Marchantia* liverworts. We compared microbiomes from *M. polymorpha*, and *M. paleacea* plants collected in the wild and their soils, all together luckily in the same geographical location (sympatric) thus reducing geographic effects; and also from plants grown in vitro and established from gemmae obtained from the same population of wild plants. Qualitative and quantitative microbiome analysis allowed us to identify microbes conserved in both native and in vitro *Marchantia* species. While *M. polymorpha* native plants microbiomes richness is reduced about M. *paleacea*, containing almost half of the Operative Taxonomic Units (OTUs) observed in *M. paleacea, M. polymorpha* grown in vitro exhibits larger OTUs. This diversity differences might be the result of impairment to recognize their microbial partners and being an open niche for opportunistic bacteria. The main OTUs in *Marchantia* microbiomes were assigned to the genera: *Methylobacterium, Rhizobium, Paenibacillus, Lysobacter, Pirellula, Steroidobacter*, and *Bryobacter*. The assigned genera correspond to bacteria capable of plant-growth promotion, complex exudates degradation, nitrogen fixation, methylotrophs, and disease-suppressive bacteria, all hosted in the relatively simple anatomy of the plant that provides refuge on their surfaces, rhizoids, and multiple gas chambers that work as specialized niches for different bacterial groups. *Marchantia* is a promising model to study not only long-term relationships between plants and their microbes but also the transgenerational impact of microbiomes because of *Marchantia* long 450 million years under climate change conditions testing microbiome configurations.

## Introduction

The evolution of life on Earth has certain milestones. Colonization of land by plants was one of the major evolutionary breakthroughs about 450 million years ago [1-4]. Land plants (embryophytes) evolved several adaptive traits that allowed them to colonize Earth’s terrestrial surface efficiently and bryophytes (mosses, hornworts, and liverworts) represent the earliest diverging land plants [5].

Evidence strongly suggest that the first land plants possessed characteristics of nonvascular plants like the *Marchantia* liverworts. *Marchantia* species are cosmopolitan, inhabits most of the planet and can colonize a high number of habitats including stream banks, rocks, logs, and as epiphytes on trees [6]. *Marchantia* includes well-known species, capable of being resistant to multiple wetting and drying cycles. Liverworts have a heteromorphic life cycle with dominant wild gametophytes [7]. *M. polymorpha* has attracted the attention of naturalists and scientists as a model system since the 18th century. *Marchantia* has been pivotal to the study of sexual chromosomes in plants, sex determination in plants in the early 1900s, the cellular nature of organisms and to the study of dorsoventral body plans and polarity in developmental biology [8,9].

Extensive parallel sequencing approaches have fostered recent studies on plant-microbe interactions, and plenty of plant microbiomes are available for the study of model plants like *Arabidopsis thaliana* [10,11]. Additionally, there are available sequenced microbiomes from plants models or with agricultural relevance. The plant microbiomes have aided in the study of plant-microbe interactions in an unprecedented way [11-15].

There are even proposed models for the establishment of the plant root microbiome like the two-step model, that considers the root microbiome as a product of plant-independent features. The two-step model considers edaphic factors, the general selection of the microorganisms for general plant cell wall composition, rhizodeposits and the host genotype which actively selects their microbial inhabitants [11]. Early land plants established symbiotic associations with mycorrhizae, and it is thought that Palaeozoic drops in CO_2_ reduced phosphate (P) absorption in nonvascular plants which in addition to competition for light favored the vascular plants Earth dominance [16].

Liverworts have a long story of endophytic relationships with fungi, and it has been suggested this is the result of a common ancient origin with multiple recent losses. For example, it has been reported that *M. paleacea* can maintain a population of endophytic fungi while some subspecies of *M. polymorpha* have lost their fungi interactor [17]. Little is known about the bacterial inhabitants of *Marchantia* other than a handful number of reports on culturable bacteria isolated from *Marchantia* thalli [18,19]. Interestingly, there are reports of *M. polymorpha* extracts able to inhibit bacterial growth (Mewari and Kumar 2008), suggesting an active microbial selection of its microbial guests. In summary, microbiome information from *Marchantia* species is practically inexistent.

In this work, we contribute the first microbiome study using massively parallel sequencing of the 16S rRNA gene for two wild *Marchantia* species, *M. polymorpha*, and *M. paleacea* along their soil substrates to inquire if there is a differential microbe selection from its surrounding habitat, considering its relevant role in plant microbiome structure. Additionally, we contribute to the general knowledge of *Marchantia* as a model plant by providing a thick description of the bacterial inhabitants of laboratory acclimated plants

## Methods

### Sampling

A total of 30 *Marchantia polymorpha* and 30 *M. paleacea* specimens were collected in wild conditions near the locality of Oxtlapa, Municipality of Xico in the Mexican state of Veracruz (19°25’26”N, 97°3’31”W; elev. 1,763 masl; collection date: 7/3/2013 3:23 p.m. UTC −6). Specimens were thoroughly collected with sterilized tweezers and contained into 50 ml sterile tubes that were frozen in place with liquid Nitrogen. Approximately 50 ml of soil volume were collected for each plant to study the role of soil as the inoculant for *Marchantia* species microbiomes. Landowners allowed us to sample within its terrain and there was no need for special collect permissions. Neither *Marchantia* species are included in the Red List of the International Union for Conservation of Nature (IUCN; http://www.iucnredlist.org/search). Gemmae from both Marchantia (female) species grown in the wild was surface disinfected and grown under *in vitro* conditions (following the detailed protocols described in Ishizaki et al. 2016) to generate thalli. We recovered gemmae from *in vitro* cultured thalli, and the gemmae obtained were grown and propagated *in vitro* for three subsequent cycles (gemmae-to-gemma). The resulting thalli were processed for DNA isolation.

### DNA extraction and sequencing library preparation

Soil and *Marchantia* species DNA was isolated using the PowerSoil DNA Isolation Kit (MoBio Laboratories, Solana Beach, CA). Briefly, five plants of each *Marchantia* species were washed with phosphate buffer as described previously [10], and the pellets were used as input for the DNA extraction. The occurring natural populations of *M. polymorpha* and *M. paleacea* were gently washed away from the surrounding particles with phosphate buffer, and then washed by vortex mixing with Tween-20 1%, the pellets were used to extract the metagenomic DNA following MoBio directions. For the *in vitro* specimens of the *Marchantia* species, the same procedure was used and the samples were removed from the agar plate with sterile tweezers, only the second wash was performed. The soil’s metagenomic DNA was extracted following MoBio’s PowerSoil indications.

The PCR primers used are the V3-V4 (341F, 805R; 464bp) region of the 16S rRNA gene (Klinsworth, 2012) which are the recommended by the MiSeq™ Illumina^®^ protocol with 5’ overhangs for multiplex library preparation:-F5’-TCGTCGGCAGCGTCAGATGTGTATAAGAGACAGCCTACGGGNGGCWGCAG-3’ and R5’-GTCTCGTGGGCTCGGAGATGTGTATAAGAGACAGGACTACHVGGGTATCTAATCC-3’. Triplicate PCR reactions were performed using a final volume of 30 μL using high fidelity Pfx Platinum polymerase (Invitrogen, Carlsbad, CA), mixing all the reactions per sample in a final volume of 90 μL. The PCR conditions are the following: denaturation at 95°C for 1 min; 5 cycles of denaturation at 94°C for 30s, annealing at 55°C for 30s, and extension at 68°C for 30s; 25 cycles of two-step cycling with denaturation at 94°C for 5s, and extension at 68°C for 30 s. PCR products were column purified using High Pure PCR Product Purification Kit (Roche Diagnostics GmbH, Mannheim, Germany).

The 16S rRNA gene sequencing libraries were constructed and sequenced at the *Unidad de Secuenciación Masiva* at the National Autonomous University of Mexico’s Biotechnology Institute, using the Illumina^®^ MiSeq™ with a 2 × 300 bp configuration according to manufacturer’s directions.

### Sequence processing

We used a previously reported pipeline [21]. Briefly, paired-end reads were merged using PANDASEQ [22], and quality control was done using fastx tools (http://hannonlab.cshl.edu/fastx_toolkit/). Briefly, all the reads were trimmed to the expected amplicon size (250 bp), then assembled using the following parameters: a minimum probability threshold of 0.95 which accounts for the minimum probability cut-off to assembly; a minimum length of 250 bp, and a maximum of 470 bp. Clustering and OTU picking was done using cd-hit-est [23] with a 97% identity cut-off over a minimum of 97% of the read length. OTU table was built using *make_otu_table.py* of QIIME’s suite [24], as well as the picking of the representative OTUs. Taxonomy assignment was conducted with BLAST against Greengenes database (13_8 release; [25]). Chimeras were identified and removed with ChimeraSlayer [26]. Finally, identified mitochondrial and chloroplast sequences were removed from following analyses.

### Diversity metrics and statistical analysis

All the diversity metrics were calculated with R (R Core Team 2014) and its phyloseq [27], and vegan [28] packages. Plots were done using R’s phyloseq, ggplot2 [29], and RColorBrewer (www.ColorBrewer.org) libraries. For the alpha diversity metrics, we calculated a number of observed species, nonparametric Chao1 index [30-32], Shannon’s index [33], and Simpson’s diversity index, for the *Marchantia* species, and for the comparative dataset. Venn diagrams were generated using the Draw Venn Diagram tool (http://bioinformatics.psb.ugent.be/webtools/Venn/). Communities distances were calculated using Bray-Curtis dissimilarities and ordered with non-metric multidimensional scaling (NMDS) with phyloseq. Several normalization procedures were conducted on the data, first for bar plots for phyla abundances relative frequency transformation was done to compare across samples/species. To compare significant differences between *Marchantia* samples, regularized logarithmic transformation (rlog) was applied to the OTUs. Counts which are computed by fitting each OTU to a baseline abundance using generalized linear model (GLM) to a baseline level, then estimating the logarithmic fold change (LFC) and dispersion for each OTU to the baseline, finally correcting for multiple testing, these comparisons were made using R’s DESeq2 package [34].

### Compared microbiomes

For comparison purposes we choose several available root microbiomes for plant species, trying to get a sparse representation of land plants. We took one *Sphagnum magellanicum* moss microbiome sequences [12]; four samples from the *Pinus muricata* microbiome study [13], here we were able to select root microbiomes from individuals with and without arbuscular mycorrhizas; two samples from B73 maize root microbiomes [14]; two samples from rice’s roots [15]; one sample from the bladder associated microbes of the carnivorous plant *Utricularia gibba* [35]; and finally, two root associated microbiomes from *Arabidopsis thaliana* and its surrounding bulk soil [10].

Sequences for similar microbiomes were downloaded from the databases declared in their respective publications. All the sequences were processed as stated before. We performed both QIIME’s *pick_closed_otus.py* and individual study OUT picking, and clustering with cd-hit-est and Greengenes taxonomy assignment for all the microbiomes and the *Marchantia* related samples.

## Results and Discussion

### Marchantia microbiome richness and diversity

*Marchantia* has emerged as an important model organism for plant developmental biology [8], and it is also a suitable evolutionary model to understand the particular adaptations of early terrestrial plants to colonize and conquer the Earth’s surface [4]. This work is the first attempt to understand the richness, and diversity of *Marchantia*’s microbial inhabitants, under wild and community established under axenic conditions [36]. The plant’s microbiome, specifically the root-associated microbiome has been shown to have dramatic effects on plants establishment, survival, and access to nutrients [11]. Given their anatomical structure, profiling microbiomes from *Marchantia*’s thalli (also containing single cell rhizoids) would be the equivalent of microbiome profiles obtained from both root and phyllosphere in vascular plants.

**Fig 1.**
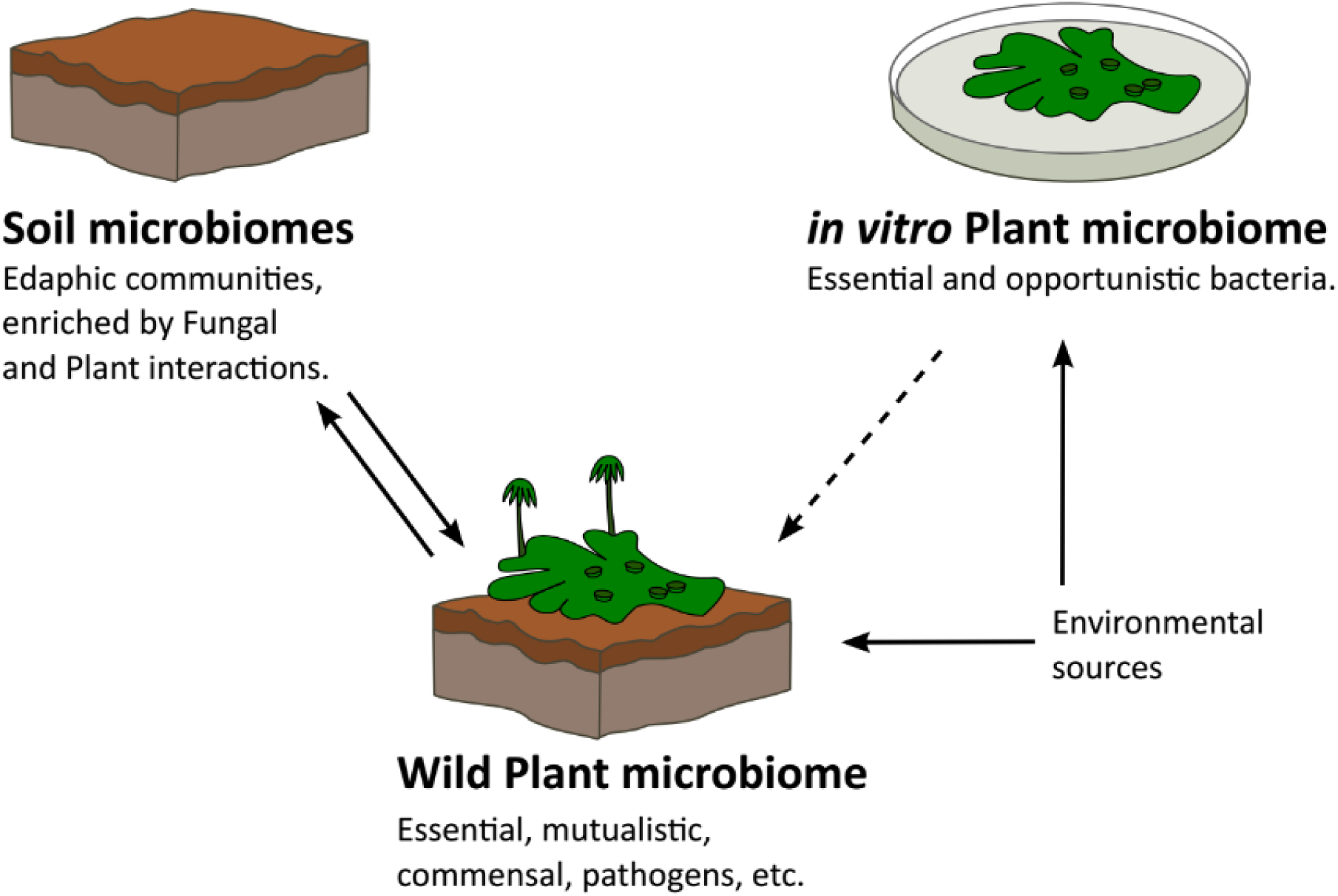
Marchantia microbiome is the product of soil microbiomes and environmental sources. Most bacteria in plant microbiome will be recruited from its soil, and soil bacteria abundances will be regulated by many fungal and plant interactions. Essential bacteria for *Marchantia* shall be present in microbiomes from plants grown in both the wild and *in vitro* conditions.

In this work, we consider wild *Marchantia* microbiomes as the result of the interactions of the plants with their environmental microbes. The soil microbiomes are the product of the interaction between their microbial inhabitants and represent a major source of microbes for plants, so we analyze the soil microbiomes for the wild *Marchantia*. Similar to other model plants, *Marchantia* can be grown in Petri dishes under supposedly axenic conditions [36]. However, there are several reports on microbe strains being isolated from *Marchantia in vitro* cultures [18,19,37]. Microbiomes of *in vitro* grown *Marchantia* are the result of careful manipulation by several generations, and it is expected that they will host less microbial diversity than a wild plant. Comparison of microbiomes from soil, wild plants and plants are grown *in vitro* aims to find bacterial inhabitants that were able to keep an intimate relationship with the plants in the restraining *in vitro* conditions and microbiomes that are the result of a long-term relationship with their *Marchantia* hosts (Fig 1).

**Table 1.** *Marchantia* OTU richness and diversity indexes. A total of 164,385 OTUs were assigned for the whole dataset, after singleton, chimeras, and contaminants removal a total of 36,299 OTUs were the base for this work. The most diverse samples are the wild *Marchantia* spp., and their origin soils. In parenthesis are shown the values of the total diversity without singleton removal. An expected dramatic decrease in diversity is observed for the *in vitro Marchantia* spp.

|  | Observed OTUs | Chao1 | Shannon | Simpson |
| --- | --- | --- | --- | --- |
| Wild *M. paleacea* | 10070<br>(17613) | 11420.56<br>(19682.06) | 8.29 | 0.998 |
| Wild *M. polymorpha* | 6188<br>(9461) | 7048.66<br>10593.97 | 7.32 | 0.996 |
| Soil *M. paleacea* | 8367<br>(15215) | 9696.03<br>(17131.73) | 8.27 | 0.999 |
| Soil *M. polymorpha* | 8313<br>(15051) | 9593.76<br>(16698.54) | 8.26 | 0.999 |
| *In vitro M. paleacea* | 402<br>(585) | 422.00<br>(603.47) | 1.35 | 0.576 |
| *In vitro M. polymorpha* | 1162<br>(1882) | 1229.50<br>(1893.86) | 3.69 | 0.903 |

A total of 10,361,954 reads were sequenced with an average of 1,726,992.33 ± 125,711.67 sequences per sample. All reads were merged into 973,945 paired-end high-quality sequences. A total of 164,385 Operational Taxonomic Units (OTUs) were assigned to the whole dataset encompassing wild populations of *M. polymorpha* and *M. paleacea*, their native soils, and the *in vitro* grown plants for both *Marchantia* species. After removal of chimeras, singletons, mitochondrial, and plant’s chloroplasts sequences, the whole dataset was reduced to 36,299 high-quality representatives OTUs for this study. *Marchantia paleacea* under wild conditions recruits almost two times more OTUs (Mpale=17,613) than wild *M. polymorpha* (Mpoly=9,461) see Table 1. Both origin soils of wild *M. paleacea* and *M. polymorpha* show even numbers in observed OTU numbers (Mpale=15,215; Mpoly=15,051). The *in vitro* grown *Marchantia* species shows an opposite pattern from its wild relatives, where *M. polymorpha* (1,882) is recruiting 3.2 more OTUs than *M. paleacea* (585). The nonparametric Chao1 index [30-32] estimate an expected number of OTUs based on the abundance of low abundance OTUs; we use the Chao1 as a reference to determine the gap between our observed OTUs and a theoretical maximum. We observed a fair sequencing coverage when comparing the observed OTUs against Chao1 Index: we have up to 89% of the wild *Marchantia* OTUs, up to 90% of soil OTUs coverage (*M. polymorpha*’s soil), and up to 99% of the *in vitro* grown *Marchantia* OTUs. Shannon’s index indicates that *Marchantia* wild species are diverse (M pale= 8.29; Mpoly=7.32), along with their soils (8.2), but the *in vitro* plants microbiomes have a substantial diversity (Mpoly=3.69; Mpale=1.35). The wild *Marchantia* spp., have large values of Simpson’s diversity index (D>0.99), the index represents the probability that two individuals sampled will belong to different species. High Simpson’s (D=0.903) stands for *in vitro M. polymorpha* (D=0.903), all but *M. paleacea* grown in the lab, which has less diversity (D=0.576).

**Fig 2.**
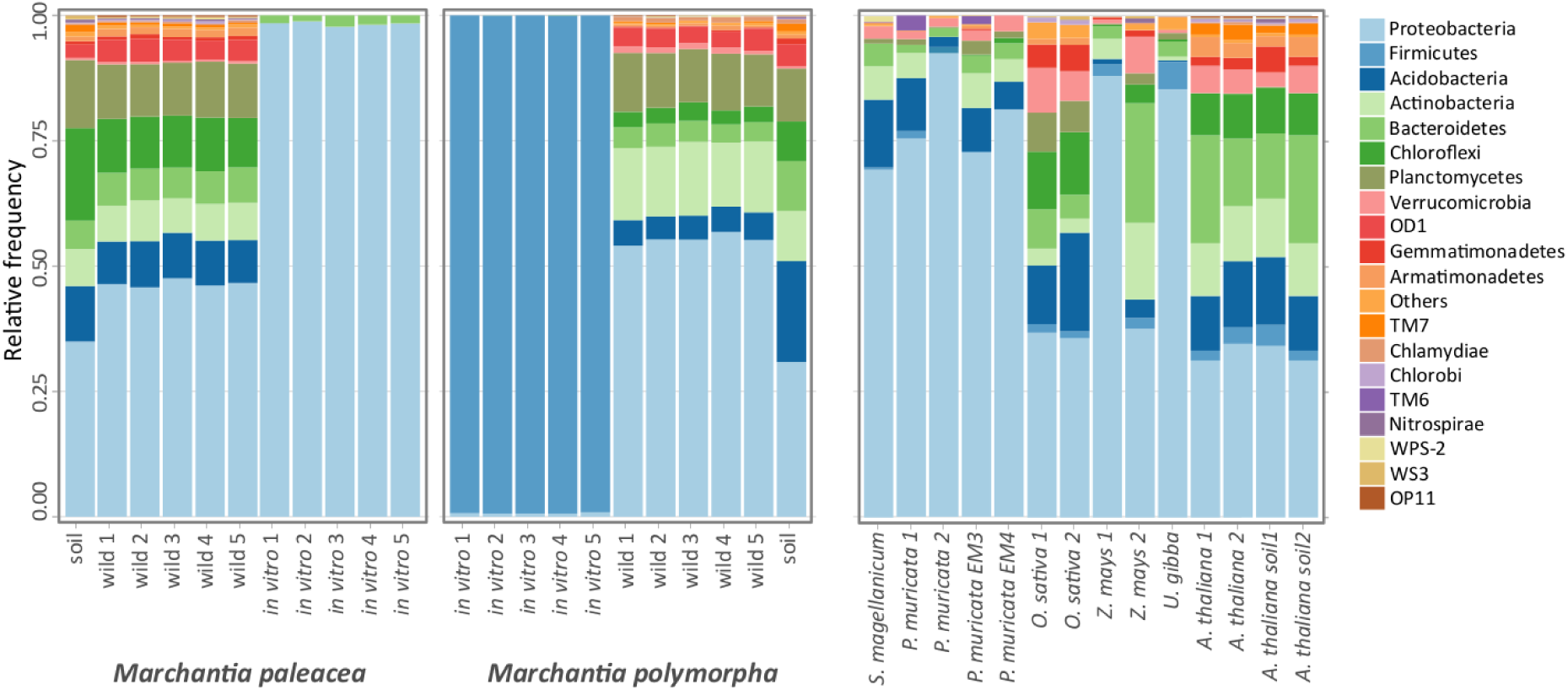
*Marchantia* microbiomes phyla diversity and a brief comparison to plant-related microbiomes. (A)*M. paleacea*, and *M. polymorpha* microbiomes. *Proteobacteria* is the most abundant phylum for both *M. paleacea* and *M. polymorpha*, all but the *in vitro M. polymorpha* with *Firmicutes* largely dominating the microbiome diversity. The two wild *Marchantia* hosts larger *Proteobacteria* OTUs than their source soils, *M. paleacea* second most abundant phylum is *Firmicutes* while in *M. polymorpha* are the *Acidobacteria.* Although there is a similar pattern in the abundance of the main phyla, the two *Marchantia* species can be distinguished based on their microbiome profiles from wild and *in vitro.* (B) When comparing to a selected set of embryophytes microbiomes, *Proteobacteria* is spotted as the most abundant phylum for every plant but in different frequencies, spanning from 30% up to 90% in the plant microbiomes. Plant microbiomes are ordered according to their phylogenetic distances: *Sphagnum* moss as a representative for bryophytes (along with *Marchantia* spp.); *Pinus* for gymnosperms; rice and maize as monocots; the carnivorous plant *Utricularia gibba*, and *Arabidopsis* to dicots (see Methods).

Taxonomic assignments of the OTUs were conducted, and here we show phyla affiliations for each of the samples, the data sets were transformed to relative frequency and are presented in Fig 2. The most abundant *Marchantia* spp. *Phylum* was *Proteobacteria* (avg=0.364) for all the samples except for *in vitro M. polymorpha* plants that host more *Bacteroidetes* (avg=0.407). In both wild *Marchantia* species the amount of *Proteobacteria*, *Bacteroidetes*, and *Verrucomicrobia* is higher in plant microbiomes than in their origin soils. On the other hand, *Planctomycetes, Chloroflexi*, and *Actinobacteria* phyla are in relative lower proportions in the wild plants about source soils (Fig 2A). The *Firmicutes* phylum is detected in all the studied samples, although they are better represented in the *M. paleacea* wild microbiome and origin soil, being barely observable in the barplot of Fig 2, neglectable in wild *M.polymorpha* and soil samples. As expected, the *in vitro Marchantia* hosted a lower overall diversity than their wild relatives being largely dominated by *Proteobacteria* and small amounts of *Bacteroidetes* for *M. paleacea*; *Firmicutes* is the main phyla, and a small fraction of *Proteobacteria* is observable the case of *M. polymorpha* (Fig 2A).

To better understand *Marchantia* microbiome results and make sense of the microbial communities described here, we performed a comparative study with other plant and soil microbiomes (Fig 2B). We compared *Marchantia* bacterial communities with microbiomes from the following plant species: The bryophyte moss *Sphagnum magellanicum*, the gymnosperm *Pinus muricata* (containing two datasets, one from roots containing arbuscular mycorrhiza and two mycorrhizae free), rice, maize, *Arabidopsis thaliana*, and *Utricularia gibba* (see Methods). By comparing the most abundant phyla across all datasets (Fig 2B), we observed common features for all the plant microbiomes such as a high relative abundance of *Proteobacteria, Acidobacteria, Firmicutes, Actinobacteria*, and *Bacteroidetes* for most of the plants analyzed. Some wild exceptions with a strong bias towards *Proteobacteria* (>0.50) are *M. polymorpha* (wild), *Pinus muricata, S. magellanicum, Utricularia gibba*, and a subgroup from the maize microbiome samples. Several non-cultivable bacteria phyla appear to be widespread distributed in plants microbiomes, albeit in lower abundances like OD1, TM7, WPS-2, WS3, and OP11. Our metagenomic DNA extraction method provided more differences in wild *Marchantia* microbiomes and their origin soils that what has been observed for *Arabidopsis* rhizospheres and soils. Phyla diversity for the *in vitro* grown *Marchantia* species we can set them apart from the rest of plant microbiomes, and *M. paleacea (in vitro)* is highly dominated by *Proteobacteria* and a remaining fraction of *Bacteroidetes.* Similar to *M. paleacea Proteobacteria* abundance is the one sample from *Pinus muricata* f203, which is an AMR infection-free rhizosphere. Interestingly, *Firmicutes highly dominate in vitro* grown *M.polymorpha* with a minimum amount of *Proteobacteria*, there is no resemblance to this pattern with any other plant/soil microbiomes analyzed. Detailed OUT tables and taxonomic assignments are available as supplementary information and illustrated as supplementary figures for the main phyla.

Using Shannon’s index we can sort the compared microbiomes based on their diversity (Table 2). Under wild conditions, *M. paleacea* hosts a larger amount of diversity than *M. polymorpha*, but this condition is reversed under *in vitro* conditions, where *M. paleacea* hosts larger diversity than *M. polymorpha*. All the compared microbiomes show large dominance values (Simpson>0.9), all but *in vitro M. polymorpha* the least dominated environment and suggesting that their microbial guests not be as structured as the other plants, and maybe the result of random opportunistic bacteria rather than of guest-host recognition and selection (Table 2).

**Table 2.** Shannon and Simpson diversity indexes for the compared microbiomes. Note that *M. paleacea* hosts larger diversity than *M. polymorpha* in wild condition but is the opposite in the *in vitro* where *M. polymorpha* hosts larger diversity than *M. paleacea*. There is a slight diversity decrease in *P. muricata* without endo-mycorrhizae.

|  | Shannon | Simpson |
| --- | --- | --- |
| *M. paleacea* wild | 8.29 | 0.998 |
| *M. polymorpha* wild | 7.32 | 0.996 |
| *M. paleacea* iv | 1.35 | 0.576 |
| *M. polymorpha* iv | 3.69 | 0.903 |
| *S. magellanicum* | 8.62 | 0.999 |
| *Pinus muricata* | 4.97 | 0.979 |
| *P. muricata* no EMR | 5.12 | 0.985 |
| *O. sativa* | 9.68 | 0.999 |
| *Z. mays* | 5.97 | 0.986 |
| *A. thaliana* | 7.19 | 0.998 |
| *U. gibba* | 8.49 | 0.999 |
| *M. paleacea* soil | 8.27 | 0.999 |
| *M. polymorpha* soil | 8.26 | 0.999 |
| *A. thaliana* soil | 7.52 | 0.999 |

**Fig 3.**
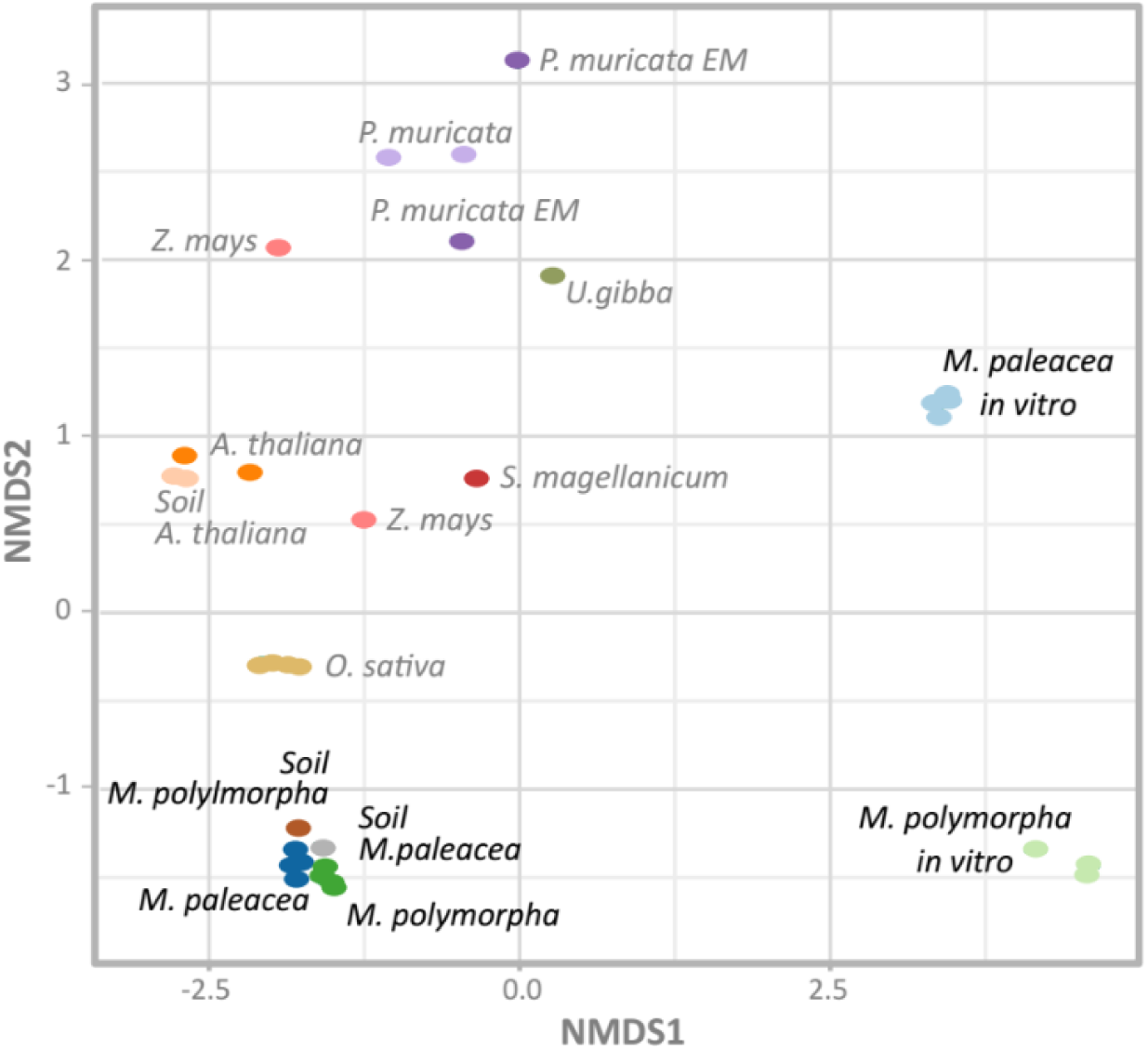
Comparative plant microbiomes ordination. The ordination was done using Bray-Curtis dissimilarities in a Non-Metric Multidimensional Scaling (NMDS). Note that *M. paleacea* and *M. polymorpha* wild plants microbiomes are clustered closer to their origin soils, and the microbiomes for the *in vitro Marchantia* are apart from every other compared microbiome. In the case of *P. muricata* microbiomes without an EM infection are grouped closer than the plants with mycorrhizal infections (NMDSstress = 0.7428).

Comparison of the microbiomes distances using NMDS and Bray-Curtis dissimilarities (Fig 3) showed that the *in vitro Marchantia* species cluster apart from every other microbiome and that they cluster together. Interestingly, there is less distance between wild *M. paleacea* and its soil, than the distance between *M. polymorpha’s* soil and microbiomes, but the two *Marchantia* species are closer in natural conditions than to any other compared plant microbiome. Additionally, the plant's microbiome tends to cluster together like is the case for rice, *Arabidopsis*, and even a sparse cluster is observed for the *Pinus*. Maize has a considerable distance between their microbiomes, but still into the left side of the NMDS1 axis, along with the plant and soil microbiomes, all but the *in vitro Marchantia* microbiomes.

A suggested explanation for the larger diversity observed in wild *M. paleacea* when comparing *M. polymorpha* is derived from previous work in which the mycorrhiza formation capabilities with glomeromycotan fungi (GA) were analyzed for a comprehensive set of species of liverworts [17]. Their work found evidence suggests an ancient symbiotic origin for GA and liverworts, as well as several independent losses for the mycorrhiza formation across the time. Ligrone and collaborators (2007) found that *M. paleacea* is capable of forming mycorrhizas with GA, while two out of three *M. polymorpha* subspecies (*polymorpha*, and *ruderalis*) were not able to form mycorrhizas with GA, while GA associations were present for *M. polymorpha* subsp. *montivagans*. They suggest that the difference between the GA presence in *M. polymorpha* could be associated with lifestyles where subspecies *montivagans* is perennial, while *polymoprha* and *ruderalis* are colonizers of nutrient rich substrates. Molecular explanations of GA absences in *M. polymorpha* could be to mutations, or gene silencing in the *M. polymorpha* mycorrhiza symbiotic genes DMI1, DMI3 and IPD3 [38]. In this work, we found a larger observed diversity both in diversity indexes and OTU numbers in the *M. paleacea* wild microbiome (17,613 OTUs), than even its soil substrate (15,215 OTUs), and almost doubles the amount of observed OTUs for *M. polymorpha* (9,461 OTUs). The lowest observed OTU number for *M. polymorpha* could be reflecting the lowest symbiotic capabilities for this plant under wild conditions. While under *in vitro* growing *M. polymorpha* has more bacteria (1,882 OTUs) than *M. paleacea* (585 OTUs). The inverse role for the plants under *in vitro* growing is possibly the result of several possibilities: 1) A cleaner microbiological condition for *M. paleacea*; 2) The microbiome of *M. polymorpha* is prone to be composed by mere opportunistic, and may be transient bacteria, while *M. paleacea* is actively recruiting its microbial inhabitants; 3) An active microbial selection and recruitment by *M. paleacea* host, which happen to be a lost trait from *M. polymorpha*; 4) Developmental timing differences between *M. paleacea* and *M. polymorpha.* Development of gemmae cups and gemmae in *M. paleacea* takes longer (~ 28 days) under our laboratory conditions. These possibilities are quite intriguing, and they do need to be explored in future works.

### Marchantia and soil microbiome interactions

**Fig 4.**
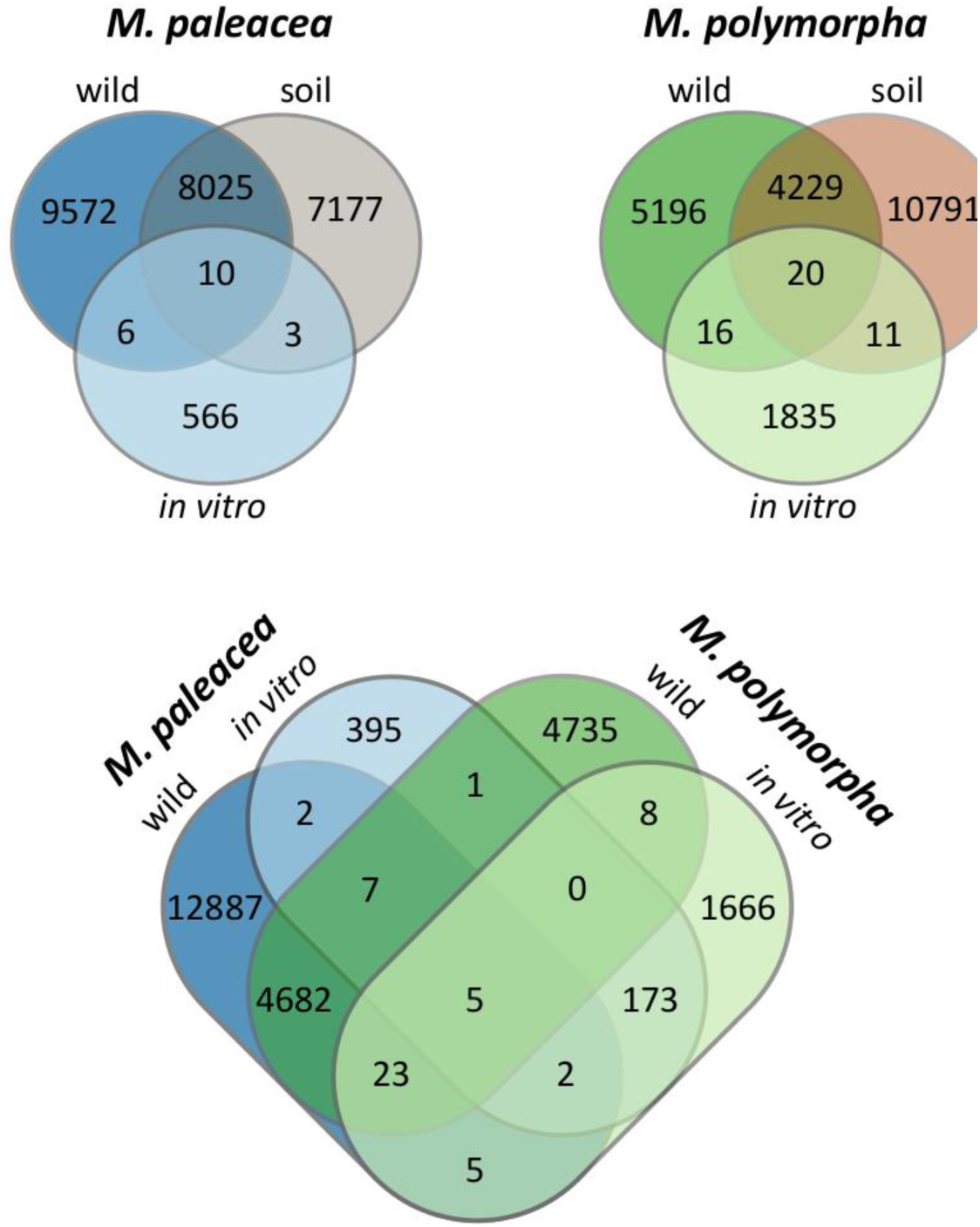
Venn diagrams of microbiomes from *M. paleacea*, *M. polymorpha*, and their soils. Above, OTUs comparison of *in vitro*, wild and soil microbiomes from each *Marchantia*. Below, microbiomes from the two *Marchantia* species from wild and *in vitro* conditions are compared.

We conducted both qualitative and quantitative analysis to determine the shared, unique, and significant differential features for *Marchantia* spp., and their soils. First, the qualitative approach using sets comparisons shows that 8,035 OTUs in *M. paleacea* are shared between *M. paleacea* and its soil; 7,180 OTUs are unique to that ground, and 9,578 are unique to the plant (Fig 4). The large number of particular plant OTUs strikes us. Some sources for these unique bacteria could be directly from air and water. However, air and water microbiomes are less diverse than soil microbial communities: in rainwater, and river microbial richness (Chao1 index) is estimated under 1,000 OTUS, while richness for air is estimated below 600 OTUS [39,40]. This diversity of microbial sources suggests that plant microbiome composition is influenced by plant-environmental interactions with some bacteria probably being inherited from plant to plant, and some bacteria pass fleetingly by soil, air, or water. There are 16 OTUs shared between wild *M. paleacea* and *in vitro M. paleacea*, these OTUs could be bacteria essential for plant growth. Another 569 OTUs are exclusive for, *in vitro* plants, they could be simply opportunistic bacteria. *M. polymorpha* case is similar; there are 4,249 shared OTUs between the wild plant and soil, and 5,212 exclusive OTUs for the wild plant. Among the wild and in vitro *M. polymorpha* just 36 OTUs are shared while 1,846 OTUs are unique to the *in vitro* plant (Fig 4).

Our analysis shows that 4,717 OTUs are common between *M. polymorpha and M. paleacea* from the wild, and 180 OTUs are in both *in vitro* grown plant species. On the other hand, the union set between the two *Marchantia* species and their soils constitutes only 5 OTUs. The identity of these five OTUs were assigned up to genus level as *Methylobacterium* (*Alphaproteobacteria*), *Rhizobium*, *Paenibacillus chondroitinus (Firmicutes)*, and the other two OTUs were (putatively) assigned as *M. organophilum. G*enera *Methylobacterium* and *Rhizobium* belong to the order *Rhizobiales*, but each one within its family.

*Methylobacterium* species have been reported as plant-associated both on phyllosphere, and roots, as forming nodules for some species, helping with nitrogen fixation, as well as cytokinin *trans*-zeatin production. They are known as pink-fermented facultative methylotrophic bacteria (PPFM), and they are capable of growing on single C sources (methanol, and methylamine), as well as other C2, C3, and C4 sources [41-43]. The possible residence of *Methylobacterium* OTUs within *Marchantia* species find support in other works were the very same *Methylobacterium* are present in year long transgenerational experiments on *Arabidopsis* [43]. Additionally, the *Methylobacterium* occurrence as a core microbiome component for the dwarf shrub *Andromeda polifolia*, which is a species of an Alpine bog vegetation which is, in turn, a community dominated by *Sphagnum* sp. [44]. Additionally, pink pigmented bacteria colonies have been observed in our *in vitro* grown *M. polymorpha*, and *M. paleacea*, and there is a report of a new *Methylobacterium* species: *M. marchantiae* sp. nov., that was isolated from a thallus of *M. polymorpha* [19]. *Methylobacterium* is a phytosymbiont for liverworts and mosses and within liverworts is proposed to consume methanol as the by-product of plant’s cell wall metabolism which is emitted through *Marchantia*’s upper epidermis stomata-like pores [19,45,46]. *Methylobacterium* is probably accumulated within plant’s air chambers*. Methylotenera* is another member of methylotrophic bacteria (C_1_ as source) over-represented in *M. paleacea*, under laboratory conditions *Methylotenera mobilis* only grows on methylamine as a source of carbon, nitrogen, and energy but under natural conditions this conditions the methylotrophy seems to be facultative [47]. This study confirms the presence of two *Methylobacterium* OTUs (147611, and 150024) that are present in soils, wild, and in vitro M. paleacea and M. polymorpha. Further work will explore the inheritance and plant’s fitness costs of the interaction with Methylobacterium and other methylotrophs.

*Paenibacillus chondroitinus* which was previously classified as *Bacillus chondroitinus* and named by its ability to degrade complex carbohydrates like chondroitin, and alginates [48] was found as a core Marchantia microbiome member. Other common soil bacteria such as *Azotobacter vinelandii* have the ability to produce alginates and is crucial for the development of desiccation resistant cysts. Bacteria produced alginates can also be used as virulence factors, like *Pseudomonas aeruginosa* which happens to infect plants, insects, nematodes, and mammals [49,50]. *Marchantia* is widely known for its desiccation resistance properties, and it is known that they can transpire the equivalent of their total water content in very short times under extreme conditions [51]. In the case of *Marchantia* accompanying bacteria, desiccation resistance strategies like the one used by *Azotobacter* (alginate mediated) or spores like the ones produced by *Bacillus* shows the importance of molecular mechanisms for drought resistance not only for the plants but their microbes. When the dry season ends it would be important to metabolize the drought resistance compounds like is the case for chondroitin. Additionally, alginate and depolymerized alginates have been shown to promote plant root growth by the induction of auxin biosynthesis genes in the plant [52,53].

**Fig 5.**
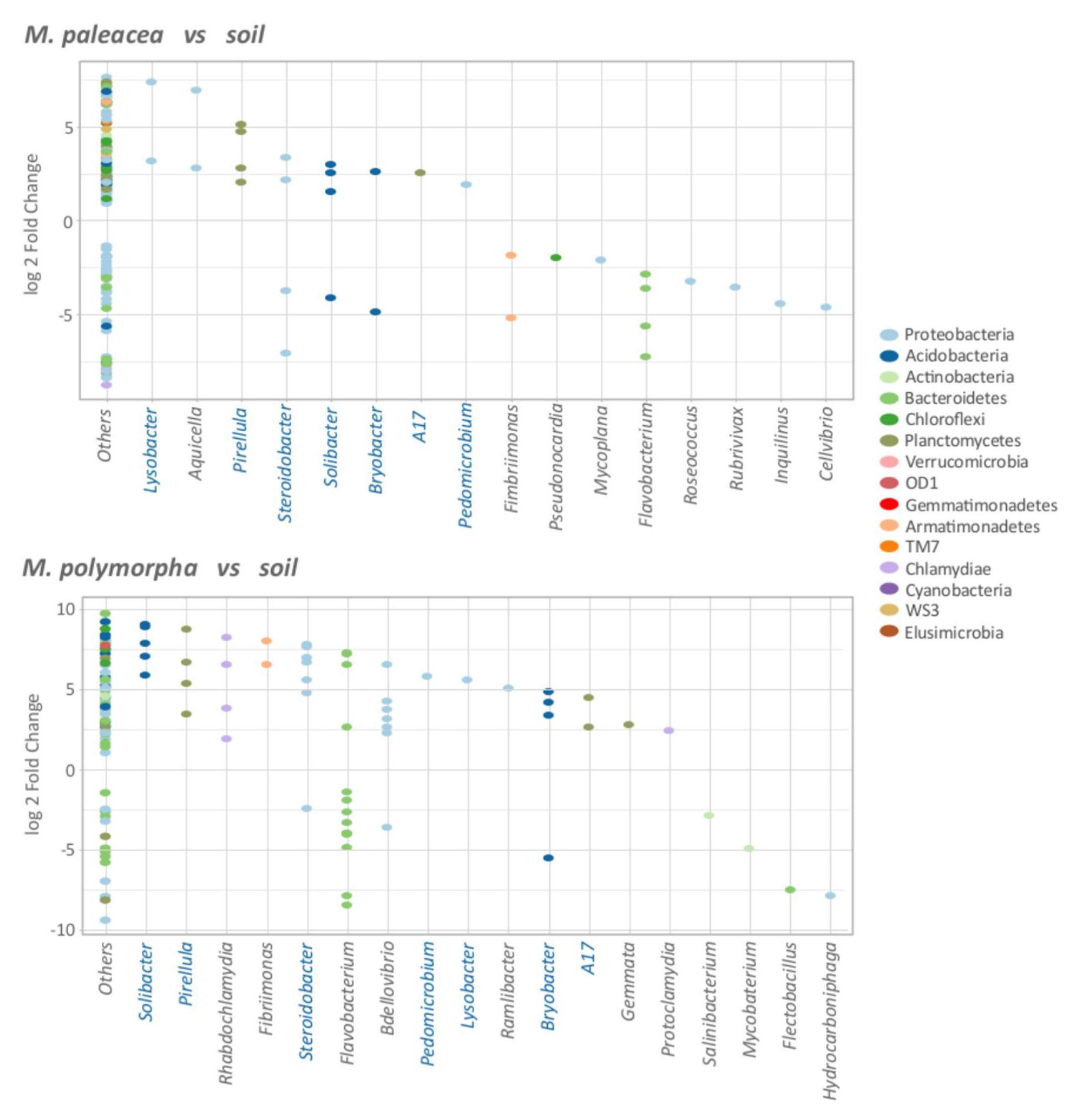
Relative abundance of the bacteria associated with *Marchantia*. The genera with significant (padj=0.01) log2 fold changes are shown. “Other” corresponds to OTUs whose genera could not be assigned. Each dot represents an OTU. The OTUs over-represented in *Marchantia spp.*, are located on top of the plot while the OTUs over-represented in soil are located at the bottom. The majority of the OTUs were not assigned to a known genus, and are shown as “Others” in the plots. Shared genera between both *Marchantia* spp., are highlighted in blue.

The quantitative approach was used to detect the strongest interactions between significant (padj=0.01) differential OTUs between plants and soil bacteria by comparing the log2 fold change (log2fc) of the OTU abundance ratio for wild *Marchantia* spp., *vs.* origin soil (Fig 5). The utility of using DESeq2 [34] log2fc where the fold changes are lower than 1 become negative values, while fold changes larger than one will be positive values, making easy to plot the changes symmetrically in a single plot. Then the comparison gets an adjusted *p-* value (padj) correcting for false positives (False Discovery Rate, FDR), using Benjamini-Hochberg (BH) correction [54]. A total of 201 OTUs of *M. paleacea* and 280 OTUs of *M. polymorpha* displayed a significant (*padj* <= 0.01) change of abundances from their soils (Fig 5). The majority of OTUs were *Proteobacteria* for both *M. paleacea* (44%), and *M. polymorpha* (35%), followed by *Bacteroidetes, Chloroflexi, Acidobacteria*, and *Planctomycetes* that are roughly around 9% (each phylum) of the significant OTUs on both plants. The “Others” category in Fig 5 correspond to the vast majority of OTUs that were not phylotyped up to genus (171/201 *M. paleacea*; 222/280 *M. polymorpha*) however they were assigned in lower taxonomical levels at least at Phylum level, and they all are bacteria with significant log fold changes due to their soils. Of the phylotyped genera, seven highly represented ones are shared in both wild *Marchantia spp.*, that could play a major role in the growth and development of the plant, these genera are *Lysobacter, Pirellula, Steroidobacter, Solibacter, Bryobacter, A17*, and *Pedomicrobium.* There are also several OTUs that are underrepresented in the plants, which indicate a negative interaction. Some genera like *Flavobacterium* or *Steroidobacter* shows OTU-specific enrichments and depletions which suggest that the relationships between plant and bacteria could be species or strain specific.

Complex carbon sources are present in plant-derived photosynthates, and thus metabolic machinery able to make use of those sources is expected to be present in their associated microbes. These specialized carbon consumers are over-represented species in wild *Marchantia* like *Bryobacter* (*Acidobacteria*) which has been described as a typical component of *Sphagnum*-dominated bogs, they are non-motile and grow well on complex heteropolysaccharides, glucuronic and galacturonic acids which are residual products of *Sphagnum* decay [55]. Other genera highly represented in *M. paleacea* are the γ*-Proteobacteria: Steroidobacter*. *Steroidobacter* has been isolated from soils and plant roots. It has been shown that some strains like *Steroidobacter agariperforans* can degrade complex polysaccharides derived from rhizospheres, and even able to degrade agar. Even more, *S. agariperforans* was isolated as a commensal strain to a *Rhizobiales* bacterium [56]. Another species, *S. denitrificans* is capable of denitrifying under anoxic conditions, using nitrate as an electron acceptor and can degrade steroidal hormones as well [57]. Recently, *Steroidobacter* has been described as part of the core rhizosphere microbiome of the gymnosperm *Welwitschia mirabilis* described as a living 110 million-year-old living fossil [58].

A nutrient rich environment like plant derived photosynthates is certainly attractive also for predators, and bacteria are no exception. *Bdellovibrio bacteriovorus* is the best studied of the *Bdellovibrio* genus; it is a sophisticated Gram-negative bacteria predator [59]. *Bdellovibrio* species has been isolated from a wide range of environments, and it is the case for plant rhizospheres. There is a report that establishes one order of magnitude major concentration of *B. bacteriovorus* in rhizospheres than circumventing soil [60] and it is very likely they are preying bacterial cells feeding on the C-rich plant exudates. *Marchantia* enrichment in *Bdellovibrio* is interesting, because it is not usually a major player in soil and rhizospheres, and it is possible it might be preying other Gram-negative inhabitants potentially pathogens as it is the case of *Agrobacterium* which happens to be over-represented in *M. polymorpha.*

Lysobacter is a Gammaproteobacteria commonly found in soils and rhizospheres. It is a bacterium with gliding motility that belongs to the family Xanthomonadaceae. Lysobacter produces antibiotics and lytic enzymes like chitinases and glucanases which protect plants from pathogenic fungi and bacteria promoting plant health and growth. Lysobacter has been proposed as a biological control agent for crop protection [61-63].

Pirellula is a Planctomycetes ubiquitous in marine and terrestrial anoxic environments. However, there are few cultivated members for the Planctomycetes. One of the few sequenced strains of formerly known Pirellula sp. 1 (Rhodopirellula baltica) revealed a potential to degrading C_1_ compounds and pathways that were thought to be exclusive for anaerobic methylotrophic bacteria and Archaea and is considered to be generating energy from the metabolism of mono or disaccharides of disaggregated algae polymers [64]. The presence of Pirellula reinforces the importance of methylotrophs within Marchantia microbiomes.

**Fig 6.**
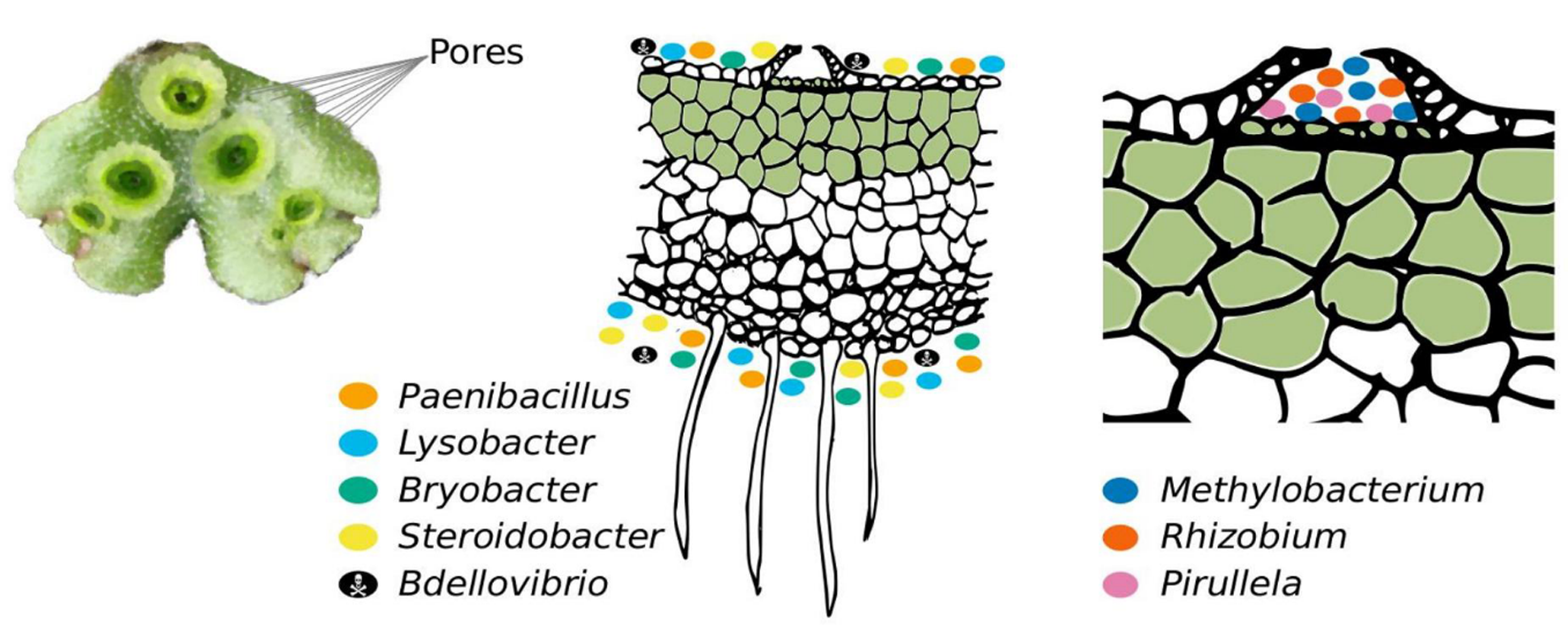
*Marchantia* microbiomes, a proposed summary. The genera described in the figure are shown as key OTUs for *Marchantia* spp. Key OTUs are proposed based on qualitative and quantitative criteria, spotlighting them as close interactors in the plants. We found genera of plant growth promoting bacteria like *Rhizobium*, and *Methylobacterium*, which have been reported as capable of Nitrogen fixing, and plant hormones synthesis. We also found complex organic compound degrading bacteria such as *Paenibacillus, Steroidobacter*, and *Lysobacter*, which have been reported to use plant-derived polymers and return plant hormones that can provide pathogen protection to their hosts. We recorded the presence of bacterial predators like *Bdellovibrio* that actively attacks and parasite other *Proteobacteria* which suggest that negative interactions among *Marchantia* spp., inhabitants might be taking place. The enrichment for methylotrophic bacteria is likely to be the result of the bacteria niche opportunity and specialization found in the *Marchantia* air chambers.

Marchantia stomata-like pores (air chambers) are visible to the naked eye and remain open as they are not fine regulated. Air chambers allow gas exchange while controlling internal moisture in contact with photosynthetic cells [51]. The importance of methylotrophic bacteria in the two Marchantia species has been highlighted as only two OTUs belonging to this genus are shared in both in vitro and wild conditions. Methanol has been reported as a normal emission of volatile organic compound (VOCs) from leaves and through plant stomata [46], and the release of plant methanol has been correlated with leaf growth [45]. The air chambers of Marchantia are a wonderful habitat for various bacteria which can make use of C_1_ sources like methanol and live within the air chambers, as it is probably the case for Methylobacterium. For both Marchantia species, we were able to identify highly abundant OTUs from their soil sources and found out that OTUs enriched in Marchantia have many pieces of evidence on related cultured strains (Methylobacterium, Rhizobium, Paenibacillus, Steroidobacter, and Lysobacter) capable of plant-derived sugar utilization, plant growth promotion, nitrogen fixation, P-solubilization. The upper taxonomic assignments for Marchantia are typical for plant derived microbiomes, and they were useful to figure out different plant microbiome distances where the in vitro grown plants cluster on their own. The microbial toolkit of M. paleacea and M. polymorpha could help us understand the microbial input to the plants and consider them in plant developmental studies. We envision some exciting avenues in research: 1) Does Marchantia’s microbiome plays a role during biotic or abiotic stress responses? 2) Is Marchantia’s microbiome transgenerationally inherited (as endophytes) or is it re-inoculated on each generation? 3) Are there differences on the microbiomes obtained from gametophytic (thalli, gametangiophores) and sporophytic tissues? 4) Can the microbiome influence Marchantia’s gene expression? 5) Can the microbiome induce epigenetic changes in the Marchantia’s genome? We believe that detailed understanding of Marchantia microbiomes and its further experimental evaluation will give us tools to efficiently test plant-microbe changes and scenarios under higher CO_2_ conditions that will be valuable for climate change modeling.

## Additional information

Accession codes: Data for the *Marchantia* microbiome has been deposited in the GenBank/EMBL/DDBJ Bioproject database under the accession code PRJNA320287. Raw genomic sequence data is deposited in the Sequence Read Archive (SRA) GenBank/EMBL/DDBJ sequence read archive under the accession code SRP078003 for the whole study and the individual accessions: SRR3746049, SRR3746050, SRR3746051, SRR3746052, SRR3746053, SRR3746054.

## Contributions

LDA, MP, AEDA and MAAV conceived the project, analyzed the data and provided reagents. LDA, MP, AEDA, JLB and MAAV wrote the paper. LDA and MAAV collected the wild soil and plants. AEDA established and perpetuated the plant material under *in vitro* conditions. MP performed metagenomic DNA extraction and library preparations for sequencing. LDA and MP elaborated the figures.

## Competing financial interests

The authors declare no competing financial interests.

## Corresponding authors

Correspondence to: Luis D. Alcaraz or Mario Arteaga

## Supporting information

S1 Fig. Acidobacteria main families.

S2 Fig. Actinobacteria main families.

S3. Fig. Bacteroidetes main families.

S4. Fig. Planctomycetes main families

S5. Fig. Proteobacteria main families.

S1 Table. *Marchantia* unfiltered microbiome OTU table

S2 Table. *Marchantia* unfiltered OTU taxonomic assignments.

S3 Table. *Marchantia* metadata file.

S4 Table. Comparative microbiomes OTU table.

S5 Table. Comparative microbiomes metadata file.

## Acknowledgements

We thank Jesús Dorantes-López, Elisa Acosta-Avilés, Liliana Arteaga-Dorantes and Elena Arteaga-Dorantes (Landowners) for permission to collect soil and plant samples. We thank Hugo R. Barajas de la Torre for sampling support. We also thank Takayuki Kohchi and Kimitsune Ishizaki for sharing unpublished protocols for *in vitro* culture of Marchantia. This research was funded by UCMEXUS Collaborative program (grant 2011-UCMEXUS-19941-44-OAC7), Consejo Nacional de Ciencia y Tecnología - Ciencia Básica (CONACYT - Basic Research) (grants CB-158550 and CB-158561), COSEAMX1 JEAI EPIMAIZE grant from the Institut de Recherche pour le Développement, Universidad Veracruzana (Cuerpo Académico CA-UVER-234), DGAPA-PAPIIT-UNAM IA200514 and CONACYT Ciencia Básica CB-237387.

